# A systems-level analysis of the mutually antagonistic roles of RKIP and BACH1 in dynamics of cancer cell plasticity

**DOI:** 10.1101/2023.07.08.547338

**Authors:** Sai Shyam, R Soundharya, Manas Sehgal, Mohit Kumar Jolly

## Abstract

Phenotypic plasticity is a hallmark of cancer metastasis. Epithelial-mesenchymal transition (EMT) is an important axis of phenotypic plasticity. Raf kinase-B inhibitor protein (RKIP) and BTB and CNC homology 1 (BACH1) are two proteins reported to influence EMT. In breast cancer, they act antagonistically, but the exact nature of their roles in mediating EMT and associated other axes of plasticity remains unclear. Here, analysing transcriptomic data, we reveal their antagonistic trends in a pan-cancer manner, in terms of association with EMT, metabolic reprogramming and immune evasion via PD-L1. Next, we developed and simulated a mechanism-based gene regulatory network that captures how RKIP and BACH1 engage in feedback loops with drivers of EMT and stemness. We found that RKIP and BACH1 belong to two separate “teams” of players – while BACH1 belonged to the one that drove pro-EMT, stem-like and therapy-resistant cell-states, RKIP is a member of a team that enables pro-epithelial, less stem-like and therapy-sensitive phenotypes. Finally, we observed that low RKIP levels and concomitant upregulated BACH1 levels associated with worse clinical outcomes in many cancer types. Together, our systems-level analysis indicates that the emergent dynamics of underlying regulatory network underlie the antagonistic patterns of RKIP and BACH1 with various axes of cancer cell plasticity, as well as with patient survival data.

## Introduction

The process of cancer metastasis is driven by phenotypic plasticity, i.e. dynamic and reversible adaptation of disseminating cancer cells to different microenvironments that they encounter along the journey. Phenotypic switching among the epithelial (E), mesenchymal (M) and hybrid E/M phenotypes through Epithelial to Mesenchymal Transition (EMT) and its reverse Mesenchymal to Epithelial Transition (MET) constitute an important axis of phenotypic plasticity during metastasis (Brabletz et al., 2018; Jia et al., 2019). Metabolic reprogramming is another key axis of plasticity and a hallmark of cancer metastasis (Celià-Terrassa and Kang, 2016). Different axes of plasticity – EMT/MET, metabolic switching and immune evasion – are often interconnected, thus enabling cancer cells to exist in distinct cell-states (Dongre et al., 2017; Jia et al., 2021; Noman et al., 2017), and promoting phenotypic plasticity and consequent non-genetic heterogeneity.

Understanding the dynamics of the interconnected axes of plasticity is critical to restrict metastasis. For instance, EMT often leads to increased levels of PD-L1 – a transmembrane molecule leading to immune escape of cancer cells (Chen et al., 2014). Further, knockdown of PD-L1 could reverse EMT (Alsuliman et al., 2015). Similarly, EMT and tamoxifen resistance in ER+ breast cancer cells can drive each other (Sahoo et al., 2021). Such bidirectional connections are often mediated by multiple feedback loops among the molecules driving cell plasticity along these multiple axes. Breaking the miR-200/ZEB1 mutually inhibitory feedback loop in breast cancer cells through CRISPR/Cas9 can reduce cancer cell metastasis (Celià-Terrassa et al., 2018). Thus, mapping the different feedback loops that can govern cell plasticity is of fundamental importance.

Recent reports have identified a mutually inhibitory feedback loop between the Raf kinase inhibitor protein (RKIP) and BTB and CNC homology 1 (BACH1) in regulation of EMT and metastasis in breast cancer (Lee et al., 2014; Wan et al., 2023). BACH1 represses RKIP transcriptionally and RKIP can inhibit BACH1 via microRNA let-7. RKIP is a metastasis suppressor that acts along the RAF1/MEK/ERK pathway to regulate cell proliferation and migration (Dong et al., 2021). Its exogenous expression in metastatic breast cancer cells can suppress invasion, intravasation and metastasis in xenograft mouse models (Dangi-Garimella et al., 2009). On the other hand, BACH1 can promote breast cancer metastasis (Yun et al., 2011). While their antagonistic roles in breast cancer have been reported, it remains unclear whether this antagonism is seen in other cancers, and how do these molecules regulate different axes of cellular plasticity implicated in metastasis.

Here, we first investigated whether RKIP and BACH1 show antagonistic trends across different cancer types using transcriptomic data from The Cancer Genome Atlas (TCGA). We found RKIP and BACH1 to be anti-correlated with each other in majority of the cancer types. Moreover, while BACH1 correlated positively with EMT and PD-L1 but negatively with oxidative phosphorylation and fatty acid oxidation, RKIP showed opposite trends in a pan-cancer manner. These trends were recapitulated in ER+ breast cancer datasets, where BACH1 also correlated negatively with ESR1 (Estrogen Receptor), but RKIP correlated positively with it. To better understand these consistent patterns in transcriptomic signatures, we constructed and simulated a mechanism-based gene regulatory network (GRN) that incorporated the feedback loops formed among RKIP, BACH1 and other master regulators of cancer cell plasticity such as ZEB1, miR-200, LIN28 and let-7. Our analysis of GRN identified that RKIP and BACH1 belonged to two mutually repressing “teams” of players – one that was comprised of pro-EMT (ZEB1, LIN28, SNAIL, SLUG) players, and the other constituted pro-MET (miR-200, let-7, miR-145, CDH1) ones. The existence of these “teams” enables the BACH1-high cells to display a hybrid E/M and mesenchymal state exhibiting a stem-like behaviour, as well as opposite trends in terms of association with patient survival across cancer types. Together, our results explain the emergent dynamics of underlying GRN that can underlie the observed antagonistic behaviour of RKIP and BACH1 in a pan-cancer manner.

## Results

### RKIP and BACH1 display mutual antagonistic patterns across many cancer types in TCGA

RKIP and BACH1 have been reported as mutually inhibitory players in breast cancer; while RKIP is anti-metastatic, BACH1 is pro-metastatic (Lee et al., 2014). To investigate whether they show antagonistic trends consistently across other cancers, we analysed TCGA data from 35 different cancer types. RKIP and BACH1 were found to be negatively correlated with each other in 31 out of 35 (88.57%) cancer types (**Fig 1A**). To understand their association with EMT, we calculated the ssGSEA score for each sample for the KS-Epithelial (KS-Epi) and KS-Mesenchymal (KS-Mes) signatures. KS-Epi signature comprises of genes that are upregulated in epithelial cells in a pan-cancer manner, while genes in KS-Mes signatures are upregulated in mesenchymal cells (Tan et al., 2014). The ssGSEA score of KS-Epi signature correlated positively with RKIP levels in only 15 cancer types and negatively with BACH1 expression in 14 cancer types, indicating a rather ambivalent association. On the other hand, the KS-Mes scores were negatively correlated with RKIP levels in 57.14% (20 out of 35) cancer types, but positively correlated with BACH1 expression in 88.57% (31 out of 35) of cases (**Fig 1A**). Further, RKIP showed a negative correlation with KS EMT score in 57.14% (20 out of 35) cancers but BACH1 showed positive correlation with it in 77.14% (27 out of 35) cases. The higher the KS EMT score, the more mesenchymal the sample is (Chakraborty et al., 2020). Together, these results suggest that RKIP and BACH1 show antagonistic trends in a pan-cancer manner, with RKIP associating with an epithelial phenotype while BACH1 with a mesenchymal phenotype.

**Figure 1:**
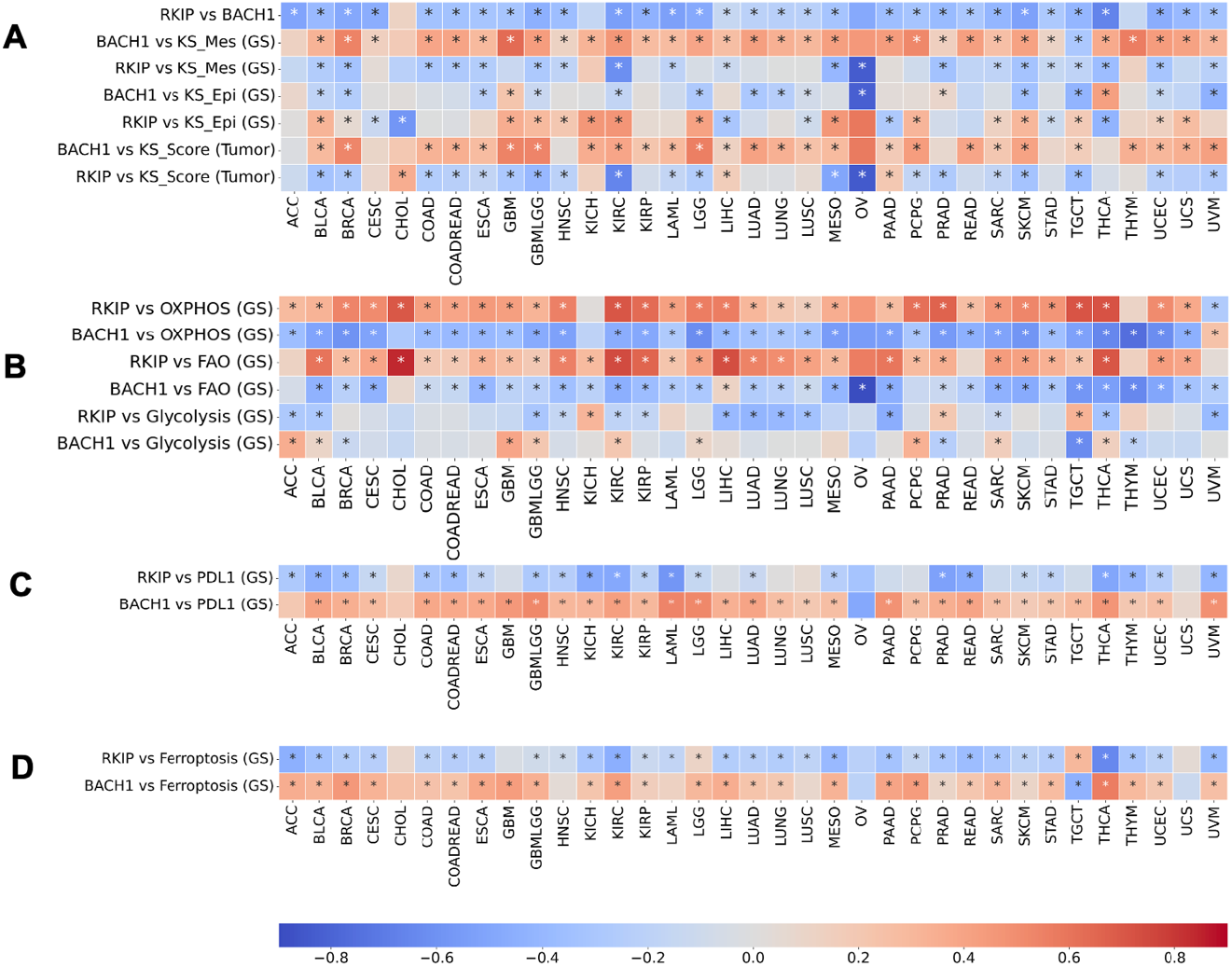
Correlation of RKIP and BACH1 expression with cellular plasticity signatures in TCGA. Heatmaps representing Spearman’s correlation coefficient of RKIP and BACH with **A)** each other, KS Score and ssGSEA scores of KS-Epi and KS-Mes gene signatures, **B)** ssGSEA scores of Hallmark gene signatures (GS) for OXPHOS, Glycolysis and FAO, **C)** ssGSEA scores for PDL1 gene signature (GS) and **D)** ssGSEA scores for ferroptosis gene signature (GS). * denotes p-value < 0.05.

During EMT, cells can also exhibit metabolic plasticity, typically leading to decreased oxidative phosphorylation and fatty acid oxidation but increased glycolysis (Muralidharan et al., 2022a). To investigate how RKIP and BACH1 expression correlates with these pathways, we calculated the ssGSEA scores for OXPHOS, FAO and glycolysis gene signatures in TCGA samples. OXPHOS ssGSEA scores correlated positively with RKIP in 88.57% (31 out of 35) of the cancer types while BACH1 correlated negatively in 91.14% (32 out of 35) of the cancer types (**Fig 1B**). Similar trends were observed with FAO, with RKIP correlating positively and BACH1 correlating negatively in 31 out of 35 cancer types. On the other hand, ssGSEA scores for glycolysis signature did not show such strong trends – they correlated negatively with RKIP in 14 cancer types, and positively with BACH1 in 9 of them. Also, while FAO and OXPHOS ssGSEA scores correlated positively with each other, glycolysis did not show consistent negative correlations with either of them (**Fig S1A)**. Overall, RKIP and BACH1 exhibited opposite patterns in terms of their association with OXPHOS and FAO.

Another axis of phenotypic plasticity coupled with EMT is immune evasion. As cells undergo a partial or complete EMT, they display increased levels of PD-L1, an immune checkpoint molecule that evades attack by the immune system (Dongre et al., 2021, 2017). Thus, we probed how the expression levels of PD-L1 and scores for PD-L1 associated gene signature correlate with RKIP and BACH1. We observed that among 35 cancer types, PD-L1 expression correlated negatively with RKIP expression in 22 of them, and positively with BACH1 in 32 of them (**Fig S1B**). Similarly, the ssGSEA scores of PD-L1 gene signature correlated negatively with RKIP in 68.57% (24 out of 35) but positively with BACH1 in 88.57% (31 out of 35) of the cancer types (**Fig 1C**). Further, cells undergoing EMT have been shown to be vulnerable to ferroptosis, an iron-dependent cell-death program (Viswanathan et al., 2017). To examine how RKIP and BACH1 associate with ferroptosis, we calculate the ssGSEA scores of a ferroptosis-based gene signature in cancer (Lu et al., 2021). Consistently, we found that RKIP associates negatively with it, but BACH1 (**Fig 1D**). Overall, BACH1 is likely associated with a more mesenchymal, glycolytic, ferroptosis-sensitive and immune-evasive phenotype, but RKIP tends to promote an epithelial, immune-sensitive and ferroptosis-insensitive cell-state dependent on OXPHOS and FAO.

### Association of RKIP and BACH1 with EMT and tamoxifen resistance in ER+ breast cancer

Next, we focused on breast cancer, given the earlier observations about mutually antagonistic roles of RKIP and BACH1 in breast cancer (Lee et al., 2019, 2014; Yesilkanal et al., 2021). We first analysed how RKIP and BACH1 correlate with *ESR1* – gene encoding for Estrogen Receptor alpha (ERα). Higher levels of ERα are often associated with improved response to anti-estrogen therapies such as tamoxifen, and better patient survival (Burns and Korach, 2012). In TCGA breast cancer samples, we observed that ESR1 correlated positively with RKIP (ρ = 0.11), but negatively with BACH1 (ρ = − 0.29) (**Fig 2A**). ZEB1, an EMT-inducing transcription factor (EMT-TF), has been shown to hypermethylated the ERα promoter and confer tamoxifen resistance (Zhang et al., 2017). Thus, we evaluated association of RKIP and BACH1 with ZEB1. ZEB1 correlated significantly positively with BACH1 (ρ = 0.52) and negatively with RKIP (ρ = − 0.32) (**Fig 2B**). Similarly, BACH1 correlated positively with another EMT-TF (SNAI2) and an EMT marker vimentin (VIM) but negatively with OVOL2, an MET-inducing transcription factor (Saxena et al., 2022), showing opposite trends as those seen for RKIP (**Fig S2A**). These results indicate that BACH1 associates with EMT, but RKIP associates with MET in breast cancer. To investigate the associations of RKIP and BACH1 with metastasis, we used their respective pathway metastasis signatures (Lee et al., 2013; Yun et al., 2011). In TCGA breast cancer samples (**Fig 2C**) and in other TCGA cancer types (**Fig S2B**), ssGSEA scores of RKIP Pathway Metastasis Signature (RPMS) correlated negatively with RKIP, while those of BACH1 Pathway Metastasis Signature (BPMS) correlated positively with BACH1. Consistently, RPMS and BPMS ssGSEA scores correlated positively with one another in a pan-cancer manner (**Fig S2B**), thus endorsing the pro-metastatic role of BACH1 and anti-metastatic role of RKIP.

**Figure 2:**
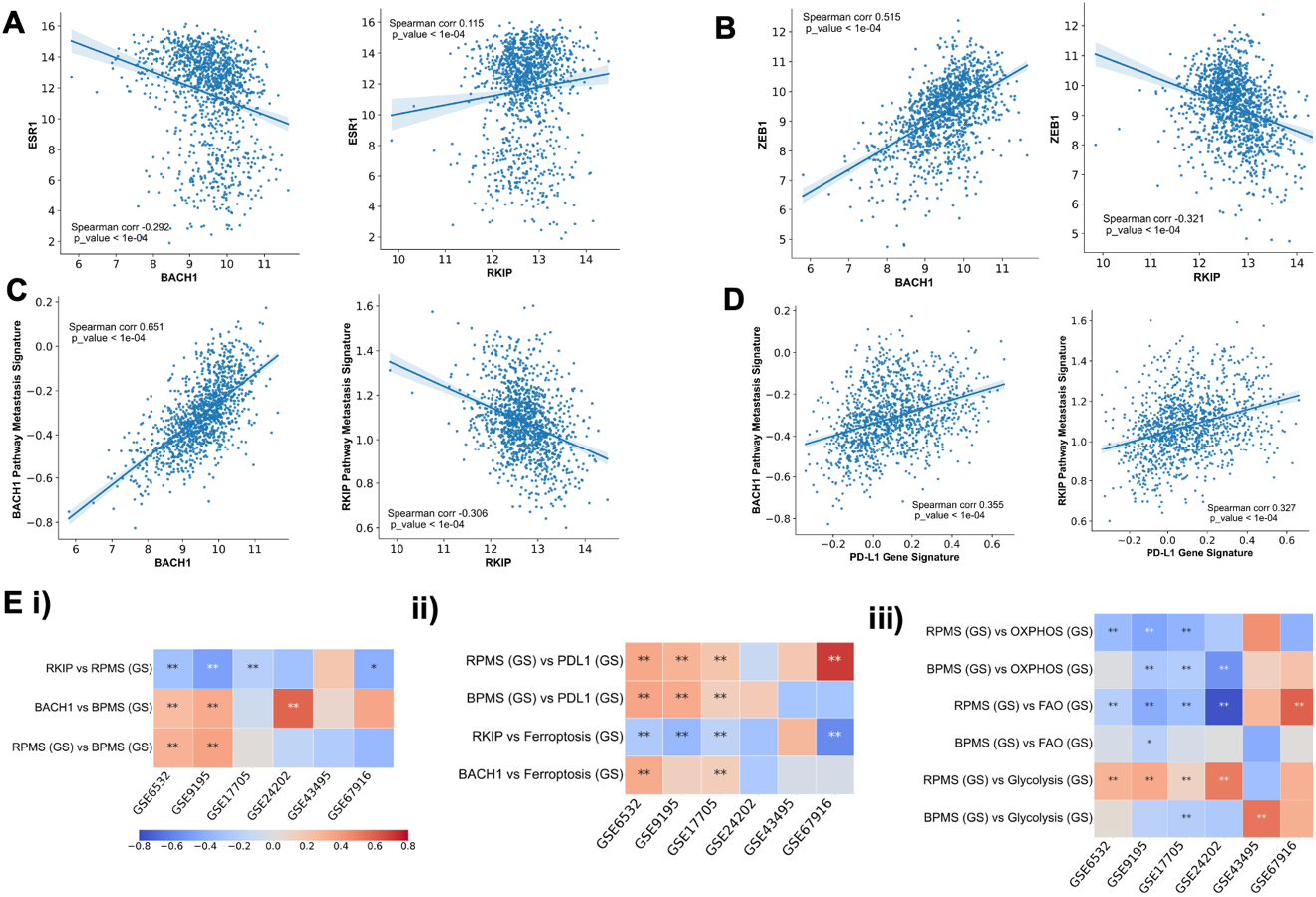
Antagonistic trends of RKIP and BACH1 with EMT & associated axes in breast cancer. **A, B)** Scatter plots showing correlations of RKIP and BACH1 with ESR1, ZEB1 in TCGA breast cancer samples. **C, D)** Same as A, B but for RKIP pathway metastasis signature (RPMS), BACH1 pathway metastasis signature (BPMS). **E)** Heatmap depicting Spearman’s correlation coefficient in 6 ER +ve Breast cancer datasets - GSE17705, GSE24202, GSE43495, GSE6532, GSE67916, GSE9195. i) Correlation of RKIP and BACH1 with BPMS, RPMS; ii) Correlation with ferroptosis and PD-L1 gene signatures (GS). iii) correlation of BPMS and RPMS with metabolic axes - OXPHOS, FAO, glycolysis. In these heatmaps, **\*\*** : p-value < 0.05, * : p-value< 0.1 (Spearman’s correlation analysis).

We next focused on six ER+ breast cancer datasets that we had previously analysed from the perspective of EMT and tamoxifen resistance driving each other in ER+ breast cancer (Sahoo et al., 2021): GSE6532, GSE9195, GSE17705, GSE24202, GSE43495 and GSE67916. We noticed resonating trends in these datasets as noted in TCGA samples: RKIP inversely correlated with ssGSEA scores of RPMS and ferroptosis gene signature, while BACH1 correlated positively with ssGSEA scores of BPMS and ferroptosis signature (**Fig 2E, i-ii**). Further, both RPMS and BPMS scores both correlate positively with those of PD-L1 signature but negatively with OXPHOS (**Fig 2E, ii-iii**), reminiscent of observations of PD-L1 activity being anti-correlated with OXPHOS across carcinomas (Muralidharan et al., 2022b). Across these datasets, RPMS correlates positively with glycolysis but negatively with FAO, while BPMS shows relatively weaker trends (**Fig 2E, iii**). This analysis further strengthens the pan-cancer trends that we noticed earlier in terms of antagonistic association of RKIP and BACH1 with multiple axes of phenotypic plasticity.

### Underlying gene regulatory networks reveal RKIP and BACH1 as members of mutually antagonistic “teams”

To understand the mechanisms that can explain the observations about RKIP and BACH1 showing opposite trends with respect to metastatic propensity, we identified a minimal core underlying gene regulatory network (GRN) in breast cancer that incorporates the feedback loops that RKIP and BACH1 are involved in with key players of EMT, tamoxifen resistance and stemness, building on our previous efforts to connect these axes of plasticity (Pasani et al., 2021; Sahoo et al., 2021). This network is not meant to be exhaustive in terms of connections RKIP and BACH1 have with players controlling these axes of plasticity, but demonstrate a core network structure that may be sufficient to explain the correlation-based observations of RKIP and BACH1 noted earlier.

Broadly speaking, this network has three core modules: EMT, tamoxifen resistance and stemness (**Fig 3A**). The EMT module comprises EMT-TFs SNAIL, SLUG and ZEB1, EMT-inhibiting micro-RNA-200 family and E-cadherin (*CDH1*), a key cell-cell adhesion molecule that maintains tight junctions among epithelial cells (Sahoo et al., 2021). The stemness module is composed of OCT4, LIN28, let-7 and miR-145. LIN28 and let-7 engage in a mutually inhibitory loop, and so do miR-145 and OCT4 (Jolly et al., 2016; Pasani et al., 2021). In tamoxifen resistance module, we include ERα66 and ERα36, two variants of estrogen receptor (*ESR1*) – ERα66 associates with tamoxifen-sensitive cell-state, while elevated levels of ERα36 drive resistance to tamoxifen in breast cancer cells (Shi et al., 2009). ERα66 can suppress ERα36 levels (Zou et al., 2009) directly, while ERα36 can inhibit ERα66 by activating ZEB1 (Sahoo et al., 2021). Other links across modules involve inhibition of LIN28 by miR-200 (Kong et al., 2010) and inhibition of ERα66 by miR-145 (Spizzo et al., 2009) as well as mutual inhibition between miR-145 and ZEB1 (Jolly et al., 2016), and that between SLUG and ERα66 (Sahoo et al., 2021a). RKIP and BACH1 associate with these modules through the following links: a) BACH1 can self-inhibit and activate SLUG (Igarashi et al., 2021), b) RKIP and SNAIL inhibit each other (Beach et al., 2007; Shvartsur et al., 2017), and c) BACH1 represses RKIP directly, while RKIP inhibits BACH1 via let-7 (Lee et al., 2014).

**Figure 3:**
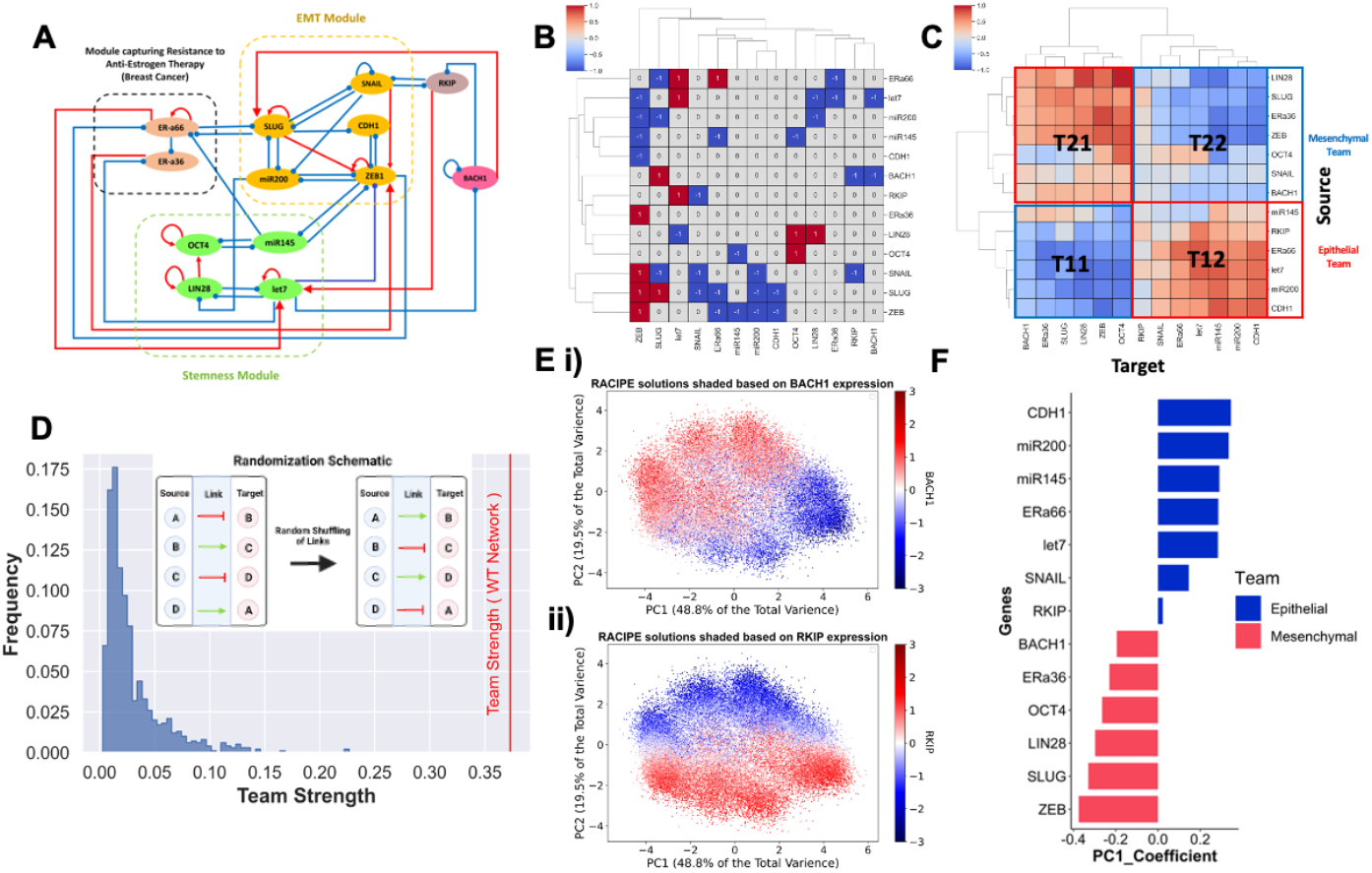
Gene regulatory network underlying antagonistic roles of RKIP and BACH1 as members of opposite “teams”. **A)** Regulatory network showing interconnections of RKIP and BACH1 with the modules of EMT, tamoxifen resistance and stemness. **B)** Adjacency Matrix for the network shown in A. **C)** Influence Matrix for path length = 10 for network shown in A. **D)** Network randomisation schematic (inset) & histogram of team strength of 1000 random networks generated. The red line indicates the team strength of the wild-type (WT) network, i.e. shown in A. **E)** Scatterplots of Principal Component 1 (capturing 48.8% variance) and Principal Component 2 (capturing 19.5% variance) shaded by BACH1 and RKIP Levels. **F)** Bar plot showing the PC1 loading coefficients of different genes in the network. The bar colors show the same antagonistic team behavior of the teams identified in C.

Before simulating the emergent dynamics of this network, we generated corresponding adjacency matrix. This matrix – showing only the direct links (both activation and inhibition) between different nodes in a network – is rather spare (**Fig 3B**). Because the influence of one node on another can also be mediated through indirect links, we derived the influence matrix for this network (Hari et al., 2022), up to a path length of eight edges, and performed hierarchical clustering (**Fig 3C**). The clustering revealed the presence of two “teams” such that members within a team effectively activated one another, while members across two teams effectively inhibited each other. One team consisted of players corresponding to an epithelial phenotype (let-7, miR-200, CDH1) while the other team comprised drivers of EMT (ZEB, SNAIL, SLUG) and stemness (LIN28, OCT4). As expected, ERα66 was found to be a part of the MET-promoting “team”, while ERα36 belonged to an EMT-driving one. Similarly, RKIP and BACH1 were found in two opposite “teams”, consistent with our previous TCGA analysis showing RKIP to be pro-epithelial but RKIP to be pro-EMT.

Next, we quantified the team strength for this influence matrix and found it to be 0.37 (on a scale of 0 to 1). To examine whether this observation of “teams” is unique to this network, we generated 1000 random networks by shuffling the edges among the nodes (following the randomization schematic; **Fig 3D** inset) and determined their corresponding team strength. We observed the team strength of the wild type (WT) network to be much greater than that of all 1000 randomly generated networks (**Fig 3D**). This observation suggests that the organization of these molecular players into “teams” in this network is not a co-incidence, rather a unique topological signature that can possibly facilitate specific couplings between cellular behaviour such as association of EMT with stemness and tamoxifen resistance (Jolly and Celia-Terrassa, 2019; Wang et al., 2019) and opposite roles of RKIP and BACH1 in EMT (Cessna et al., 2022; Igarashi et al., 2021).

Further, we simulated the dynamics of this GRN by representing the regulatory interactions through a set of coupled ordinary differential equations (ODEs) across an ensemble of kinetic parameters and initial conditions, using a tool called RACIPE (Huang et al., 2018). The output of RACIPE is the set of steady-state solutions obtained which are then z-normalized for a better comparison across the expression patterns. Principal component analysis (PCA) performed on RACIPE solutions indicated that over 60% of the variance could be captured by first 2 principal components (PCs) (**Fig S3A**) - PC1 accounted for 48.8%, while PC2 accounted for 19.5%. Colouring the PCA plots based on z-normalized levels of RKIP and BACH1 revealed that clusters showing higher BACH1 levels had lower RKIP levels and *vice versa* (**Fig 3E**). Next, we plotted the loading coefficients of each node in the network along the PC1 axis to understand their contributions to PC1 variance. We observed that genes earlier identified to be a part of the epithelial team from the influence matrix analysis had PC1 coefficients less than zero, while all of them in mesenchymal team (with SNAIL being the only exception) had these coefficients greater than zero. This analysis reveals that PC1 is largely able to recapitulate the members belonging to two different “teams” identified via influence matrix. The existence and constitution of these “teams” is further validated by largely bimodal distribution of all network nodes, PCA correlation circle and pairwise correlation matrices showcasing positive correlation among members within a team, and negative across “teams” (**Fig S3B-C**).

### Regulatory network dynamics explains the association of BACH1 with stem-like cell-state

Next, we investigated whether this regulatory network is capable of multi-stable behaviour, i.e. allowing for the co-existence of multiple cell-states that can reversibly switch among themselves. We segregated the parameter sets generated by RACIPE based on their corresponding number of stable states (**Fig 4A**). Our analysis revealed that only 4.58% of solutions correspond to a mono-stable state; the remaining 95.42% solutions associated with two or more co-existing cell-states, highlighting the underlying multistable dynamics of this GRN that can support phenotypic plasticity.

**Figure 4:**
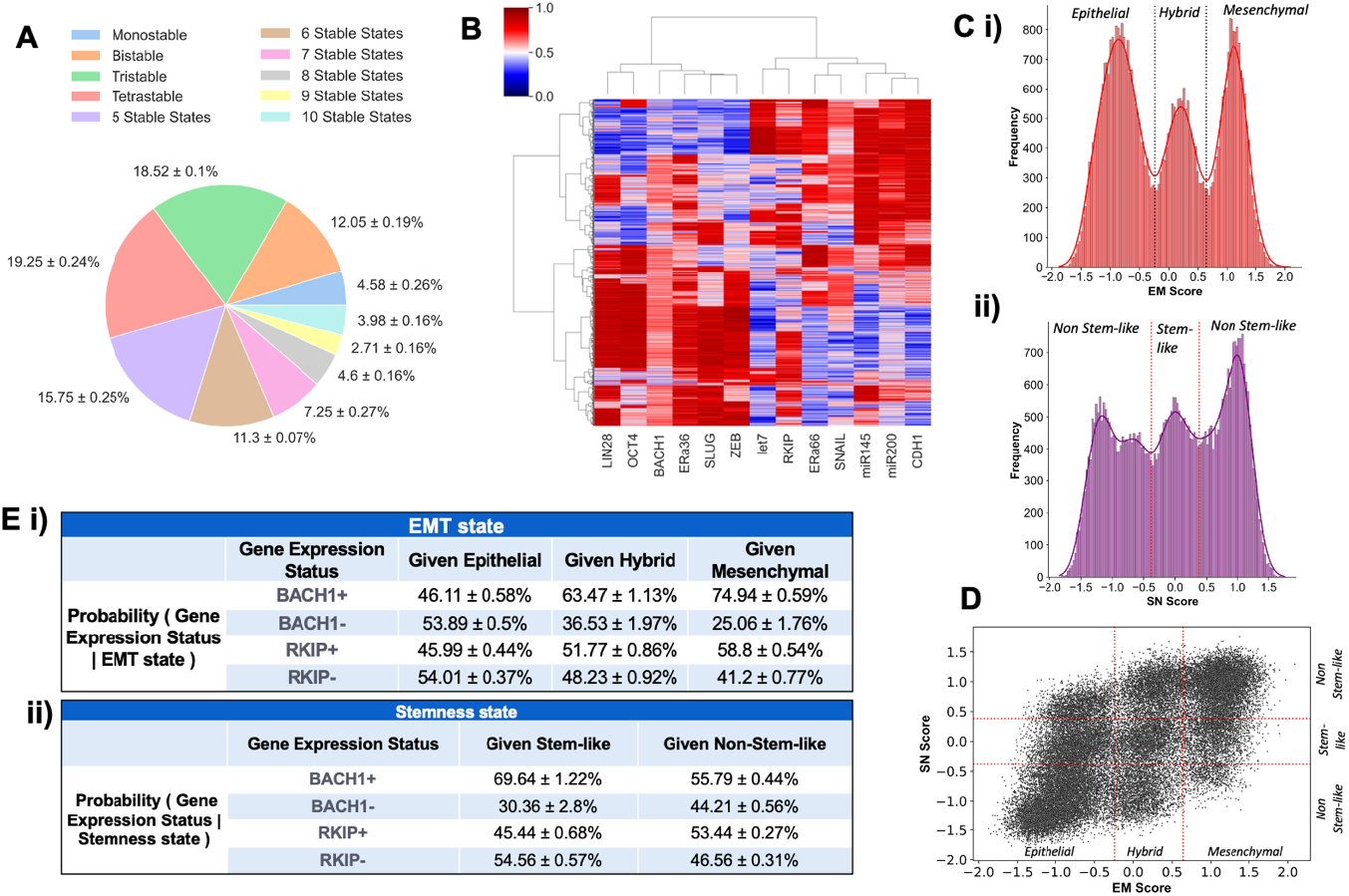
Association of RKIP and BACH1 with EMT and stemness states. **A)** Pie chart depicting the number of parameter sets giving rise to different number of stable states. Data from 3 independent replicate RACIPE simulations shown as mean ± standard deviation). **B)** Heatmap of stable steady-state solutions for network shown in Fig 3A, simulated from RACIPE. Color bar represents the relative levels of individual genes (z-normalized and log2 expression values). **C)** i) Kernel Density Plot of EM Score (= ZEB +SLUG – miR200 – CDH1). Red dotted lines indicate the segregation of Epithelial, Hybrid and Mesenchymal phenotypes based on minima of the fitted Gaussians. ii) Kernel Density Plot of SN Score (= OCT4 +LIN28 – mir145 – let7). Red dotted lines indicate the segregation of stem-like and non-stem-like phenotypes based on minima of the fitted Gaussians. **D)** Scatterplot of all RACIPE solutions, on SN Score vs EM Score axes. Segregation done based on thresholds identified in C i, ii. **E)** i) Table depicting the conditional probability percentage of gene expression status given the epithelial, mesenchymal or hybrid Phenotypes. ii) Same as i) but given the stem-like and non-stem-like categorization. BACH1 (or RKIP) +ve indicates z-score expression of BACH1 (or RKIP) > 0, and BACH1 –ve indicates it < 0.

Further, we plotted the ensemble of stable states obtained from RACIPE as a heatmap (**Fig 4B**). It showed expected clustering patterns and the dominance of two distinct states – (high LIN28, high OCT4, high BACH1, high ERα36, high SLUG, high ZEB, low let-7, low ERα66, low SNAIL, low miR-145, low miR-200, low CDH1) and (low LIN28, low OCT4, low BACH1, low ERα36, low SLUG, low ZEB, high let-7, high ERα66, high SNAIL, high miR-145, high miR-200, high CDH1), thus recapitulating the “teams” seen in influence matrix of the GRN. These two states correspond to a mesenchymal, stem-like, tamoxifen-resistant and epithelial, non-stem-like, tamoxifen-sensitive state respectively. Interestingly, higher levels of RKIP were seen in most RACIPE solutions that correspond to an epithelial cell-state, as well as a subpopulation of mesenchymal cell-state subpopulation. This observation is consistent with a lower magnitude of loading coefficient of RKIP in explaining PC1 variance as compared to that of BACH1 (**Fig 3D**). Thus, our simulations suggest that BACH1 associates strongly with a partial/full EMT state, relative to the association of RKIP with an epithelial one.

To better elucidate the functional mapping of phenotypes from the RACIPE expression data, we defined an Epithelial Mesenchymal (EM) score as EM Score = (ZEB1 + SLUG − miR200 − CDH1)/4. ZEB1 and SLUG are characteristic EMT markers, while miR-200 and CDH1 are those for an epithelial state. The higher the EM score, the more mesenchymal the corresponding cell-state is. A histogram of EM scores thus obtained across all RACIPE solutions indicated three distinct populations which can be characterized as epithelial, mesenchymal and hybrid ones (**Fig 4C,i**). Similarly, to quantify plasticity along the stemness axes, we defined a stemness (SN) score as SN score = (OCT4 + LIN28 − miR145 − let7)/4. Extremely high or low levels of OCT4 and LIN28 are associated with non-stem-like states (Jolly et al., 2014; Karwacki-Neisius et al., 2013; Niwa et al., 2000), thus we ascribe intermediate levels of SN scores to a stem-like state, while very high or low levels of it to a non-stem-like state, based on the histogram (**Fig 4C,ii**).

The z-normalized RACIPE solutions were then projected on a scatter plot with corresponding SN and EM scores as the axes. While the scatter plot showed a general positive correlation between EM and SN scores (ρ = 0.701), we noticed associations of all the three phenotypes along the EMT axis with both stem-like and non-stem-like states (**Fig 4D**). To better understand how RKIP and BACH1 levels associated with these axes, we divided the RACIPE steady state solution ensemble into BACH1 (+ve) and BACH1 (-ve) (and similarly for RKIP (+ve) and RKIP (-ve)) cases based on their corresponding z-normalized values. We observed that 63.48% of *in silico* cells (considering each steady-state solution of RACIPE as equivalent to a cell in an experimental population-level setting) showing a hybrid E/M phenotype were BACH1(+ve) (**Fig 4E,i**). Similarly, a mesenchymal cell was three times more likely to be BACH1(+ve) as compared to being BACH1(-ve) (74.94% vs 25.06% cases) (**Fig 4E,i**). Consistent with experimental observations of partial and/or full EMT phenotypes with stemness (Brown et al., 2022; Lourenco et al., 2020; Lüönd et al., 2021), we found a stem-like cell to be 2.3 times (69.64% vs 30.36%) more likely to be BACH1(+ve) as compared to being BACH1 (-ve) (**Fig 4E, ii**). On the other hand, the RKIP(+ve) or RKIP(-ve) cells were not enriched in any subpopulation along the EMT or stemness axes. Together, we can infer that BACH1 associates with a hybrid E/M and/or mesenchymal stem-like phenotype.

### High BACH1 expression levels associate with worse patient survival in many cancer types

To identify how RKIP and BACH1 expression levels impact survival outcome, we computed the hazard ratios for different combinations of RKIP and BACH1 (RKIP+ BACH1-, RKIP+ BACH1+, RKIP-BACH1-) with RKIP-BACH1+ as reference. High BACH1 expression along with low RKIP expression led to significantly worse overall survival when compared to high RKIP and low BACH1 samples in three different cancers: lung adenocarcinoma (LUAD), liver hepatocellular carcinoma (LIHC) and pancreatic adenocarcinoma (PAAD). Similar trends were recapitulated for disease specific survival, disease-free survival and progression-free survival in these cancers. Conversely, RKIP+ BACH1+ samples had significantly better progression free survival compared to the RKIP-BACH1+ in LIHC, indicating that higher RKIP expression may be linked to better survival outcomes (**Fig 5**). Consistent trends are also observed in mesothelioma, breast cancer, renal clear cell carcinoma, uveal melanoma and thyroid carcinoma (**Fig S4**). These pan-cancer observations carried out consistently indicate that high BACH1 levels are linked to worse survival.

**Figure 5:**
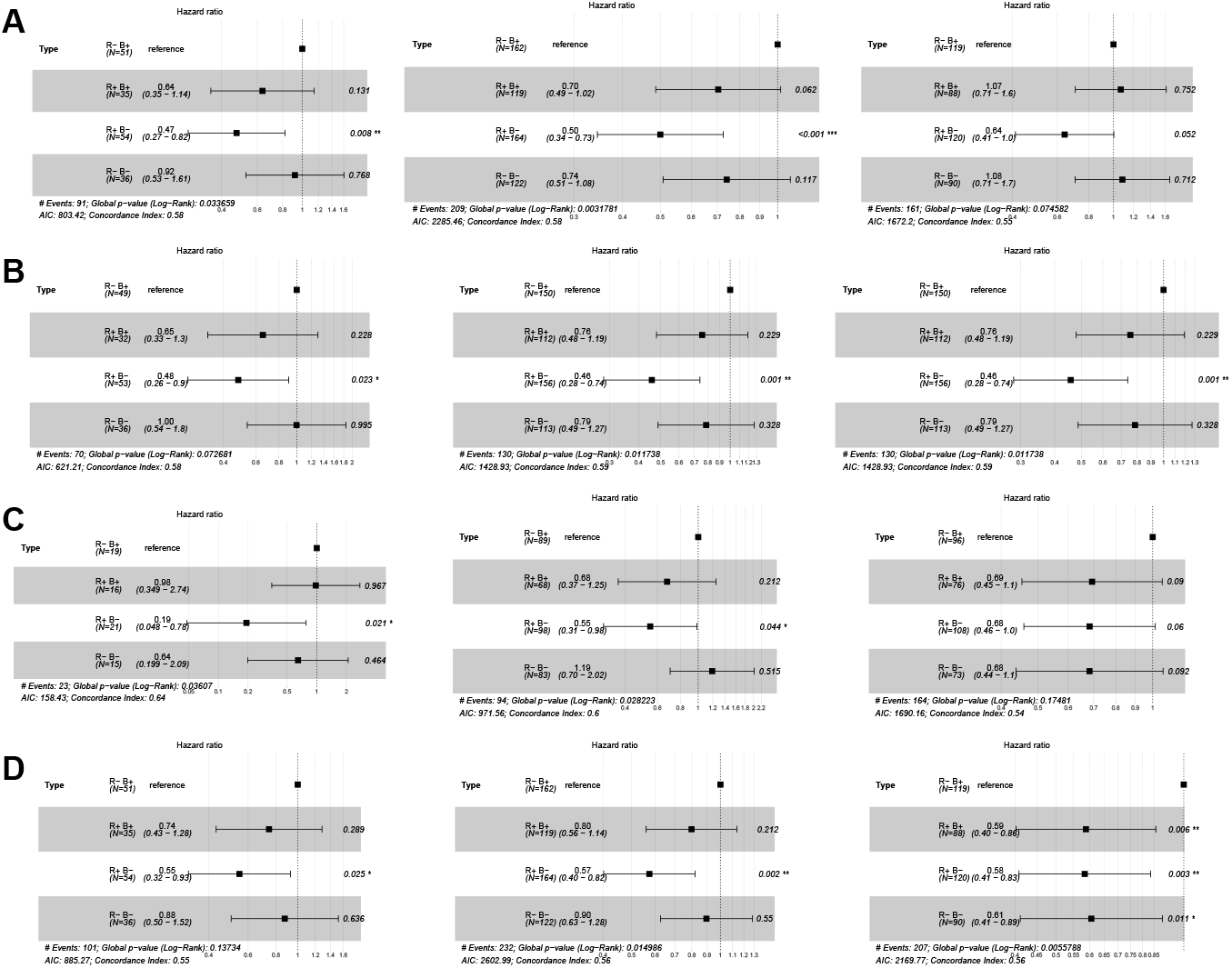
Forest plots showing antagonistic trends of RKIP and BACH1 with varying survival types across different cancers. Reference set of TCGA sample considered here is R-B+ (RKIP-low, BACH1-high). p-values are based on log-rank test, and those with significant differences p < 0.05, p<0.01 and p<0.001 are marked with *, **, and ***, respectively. **A)** Hazard Ratios (HR) for Overall Survival in PAAD (left), LUAD (middle) and LIHC (right). **B)** HR for Disease Specific Survival in PAAD (left), LUAD (middle) and LIHC (right). **C)** HR for Disease-Free Survival in in PAAD (left), LUAD (middle) and LIHC (right). **D)** HR for Progression -Free Survival in PAAD (left), LUAD (middle) and LIHC (right).

## Discussion

Cellular decision-making is often governed by mutually inhibitory feedback loops, also called as a ‘toggle switch’. Such network motifs are commonly reported in various decision-making contexts, such as GATA1-PU.1 toggle switch determining differentiation of a common myeloid progenitor into erythroid cells (GATA1 >> PU.1) or myeloid cells (PU.1 >> GATA1) (Zhou and Huang, 2011). Similar antagonism has been witnessed among many EMT and MET driving transcription factors (TFs) such that the MET-TFs (GRHL2, KLF4, OVOL2 etc.) and EMT-TFs (ZEB1, SNAIL, TWIST etc.) mutually repress one another (Chung et al., 2016; Roca et al., 2013; Subbalakshmi et al., 2021). Such mutually antagonistic “teams” of players – also seen in small cell lung cancer GRN (Chauhan et al., 2021) – can thus contribute to cellular plasticity (Hari et al., 2022).

Here, we show that RKIP and BACH1 belong to two antagonistic “teams” – while RKIP belongs to a “team” promoting an epithelial and tamoxifen-sensitive cell-state in breast cancer, BACH1 is a part of “team” enabling a mesenchymal and tamoxifen-resistant phenotype. While the antagonistic role of RKIP and BACH1 in EMT has been well-reported, their potential involvement in tamoxifen resistance is only beginning to be investigated through mediators such as TANK-binding kinase 1 (TBK1) (Liu et al., 2022; Wei et al., 2014). Consistently, their contrasting roles are emerging in the contexts of ferroptosis (Wenzel et al., 2017; Xie et al., 2023) and metabolic reprogramming (Lee et al., 2019) in specific cancer types. Particularly, activation of NRF2 – a master regulator of anti-oxidant program in cells – stabilizes BACH1 (Lignitto et al., 2019). NRF2 activity can be enhanced by loss of RKIP (Al-Mulla et al., 2012), enabling another ‘toggle switch’ between RKIP and BACH1. NRF2 has also been reported to maintain cells in a hybrid epithelial/mesenchymal (E/M) phenotype (Vilchez Mercedes et al., 2022) and enhance stemness and chemoresistance (Al-Mulla et al., 2012; Kim et al., 2018). Thus, our results about association of BACH1 with a hybrid E/M stem-like state unify the previous observations about role of BACH1 in controlling multiple axes of plasticity.

The contrasting roles of RKIP and BACH1 in mediating stemness/dedifferentiation lends further credence to our model simulations. BACH1 can activate CD44 and MAPK signaling in lung cancer stem cells (CSCs) and stimulate lung cancer metastasis; its loss represses metastasis in xenograft models (Jiang et al., 2021; Wiel et al., 2019). Similarly, BACH1 represses mesendodermal differentiation in embryonic stem cells, maintaining stem-cell identity (Wei et al., 2019). Conversely, RKIP can repress NANOG in primary melanocytes, maintaining their differentiation state (Penas et al., 2020). Chemical induction of RKIP can degrade SOX2, inhibit tumor growth and promote differentiation of schwannoma into mature Schwann cells (Cho et al., 2022). These observations, together with the contrasting roles of RKIP and BACH1 on tumor cell migration (Davudian et al., 2016; Zhu et al., 2018), can explain the association of higher BACH1 levels with enhanced metastasis and poor patient prognosis (Chen et al., 2023; Han et al., 2019; Wiel et al., 2019). Our results demonstrate that these trends of association of RKIP and BACH1 with clinical outcomes are largely consistent across cancer types as well as survival metrics (overall survival, progression-free survival, disease-free survival), highlighting their behavior as members of two “teams” playing a tug-of-war – a pro-metastatic one (EMT, stem-like, drug-resistant) and an anti-metastatic one.

Overall, our pan-cancer systems-level analysis reveals that RKIP and BACH1 can control multiple axes of plasticity (EMT, metabolic reprogramming, stemness) together in opposite directions, thus explaining their association with patient survival seen across cancer types. Our mechanistic model highlights that such “teams” of players can be an important network motif in co-ordinating changes along more than one axes of plasticity together, such as EMT, stemness and metabolic switching. Our results suggest breaking such “teams” as a possible therapeutic avenue to reduce the fitness of metastasizing cells, by limiting their phenotypic plasticity trajectories.

## Conflict of Interest

The authors declare no conflict of interest.

## Author contributions

SS, SR and MS performed research and analysed data. SS prepared initial draft of the article, and all authors edited it. MKJ designed and supervised research and acquired funding.

## Acknowledgements

This work was supported by Ramanujan Fellowship (SB/S2/RJN-049/2018) awarded to MKJ by Science and Engineering Research Board (SERB), Department of Science and Technology, Government of India.

## Code and Data Availability

https://github.com/saishyam1/RKIP_BACH1_Data_Codes.git

## Materials & Methods

### 1. Transcriptomic datasets

We downloaded microarray data from NCBI GEO (GSE17705, GSE24202, GSE43495, GSE6532, GSE67916, GSE9195) using the GEOquery package on R, and mapped the probes to respective genes using their corresponding annotation files. We normalized gene-wise expression matrices on log2 base, before further analysis.

We downloaded TCGA gene expression data of 35 different cancer types, obtained from the UCSCXena browser. These pre-processed datasets were available in transcripts per million (TPM) format and were directly used for analysis. Survival data was analysed using data from TCGA. Relevant cancer samples were split into two groups based on median: high *PEBP1* (gene name for RKIP) vs low PEBP1, and high *BACH1* vs low BACH1. ‘Kaplan -Meier curves for overall survival were plotted using the plotter on the ‘KMPlotter’ website (Nagy et al., 2021). Additionally, ‘coxph’ function in R package ‘survival’ was employed to determine the hazard ratio (HR) and confidence interval (95% CI) for TCGA cohorts, and heatmaps were made using ‘ggplot2’.

Normalized Single Sample Gene Set Enrichment Analysis (ssGSEA) scores based on specific input gene signatures (**Table S1**) were calculated for each sample using ‘GSEApy’ python package (Subramanian et al., 2005). For correlation analysis between any two variables, Spearman’s correlation coefficient has been used, using ‘spearmanr’ function from ‘SciPy’ Python library.

### 2. EMT scoring metric

KS score is a metric to quantify the extent of EMT based on expression levels of specific epithelial and mesenchymal markers (Tan et al., 2014). It uses 2 gene signatures: KS-Epi (genes associated with an epithelial phenotype), KS-Mes (genes associated with a mesenchymal phenotype). It plots two cumulative distribution functions (CDFs) based on expression levels of genes in KS-Epi and KS-Mes signatures. Distance between the 2 CDFs is calculated for each value, and the maximum value is taken as the statistic. The KS score values lie in the range [-1, 1].

### 3. Identification of “teams” of players

An adjacency matrix (Adj) represents the topology of a GRN in the form of a matrix. Rows represent the source node for a particular link, while the columns represent the target nodes. An inhibitory link is represented (in blue) with value -1; an activation link is represented (in red) with value 1. Path-length is defined as the number of consecutive links in a path that connects a source node to its corresponding target. In a GRN, the nodes not only influence their direct target (path length = 1) but also other nodes indirectly (path length > 1). To take into consideration these interactions, we define an influence matrix that captures both the direct and indirect interactions between nodes in the network up to a defined path length (Hari et al., 2022).

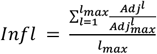

where Adj^l^ represents the adjacency matrix (Adj) multiplied with itself l times. Adj_max_ represents the Adj matrix with all inhibition links replaced by activation links. Adj^l^ / Adj^l^_max_ represents the element-wise division of values in Adj^l^ by values in Adj^l^_max_.

A positive value indicates activation, and a negative influence indicates inhibition. The higher the value in the influence matrix, the higher the influence of that specific source gene on the target.

Hierarchical clustering is performed on the influence matrix to identify the clusters of genes functioning similarly. The team Strength of each cluster T_KL_ is given as

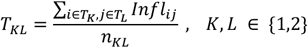

Team Strength of the entire network T_S_ is given as

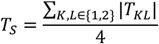

### 4. RACIPE analysis

Random Circuit Perturbation (RACIPE) is a tool used to simulate the dynamics of GRNs. It takes as input the topology of a GRN and generates an ensemble of kinetic models for the given GRN. Here, we simulated the network shown in Fig 3A – the input file to RACIPE is given as **Table S2**.

For each kinetic model generated for the input topology file, RACIPE samples many initial parameters from the designated range for each parameter. The expression levels of a node in a GRN is determined by a set of ordinary differential equations given below.

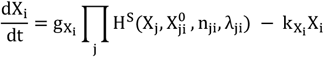

Here, X_i_ is concentration of gene product of gene node X part of GRN and 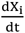 is the rate of change of the gene expression with respect to time. 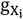 is basal production rate; 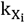 is basal degradation rate of gene product X_i_. H^s^ represents shifted Hills function that models the activation and inhibition links towards this (X_i_) gene node in the GRN. n_ji_ is Hill’s function coefficient and 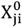 is threshold value of the Hill equation. X_j_ is concentration of gene product X_j_, where X_j_ is a node either activating or inhibiting X_i_. Since our GRN had 13 nodes, i and j can take integer values between 1 and 13. λ_ji_ is fold change parameter. Throughout this study, we simulated GRNs for 10,000 parameter sets. RACIPE gives log_2_ normalized steady state gene expressions of each gene product as output. This output steady state gene expression data was then z-normalized and used for analysis. To check for bimodality in solutions obtained from RACIPE, we used Sarle’s bimodality coefficient:

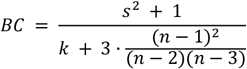

Here, ‘s’ is the skew of the distribution and ‘k’ is the kurtosis. BC can take values between 0 and 1, with BC values greater than 0.55 indicating bimodal distribution of gene expression (Sahoo et al., 2020). SciPy library has been used to calculate kurtosis (k) and skew (s) of the distribution.

## Supplementary Table Legends

Table S1 – List of gene signatures used in ssGSEA, along with their corresponding reference.

Table S2 – List of network nodes and edges for the network in Fig 3A, given as an input topology file to RACIPE, along with their corresponding reference.

## Supplementary Figures

**Figure S1:**
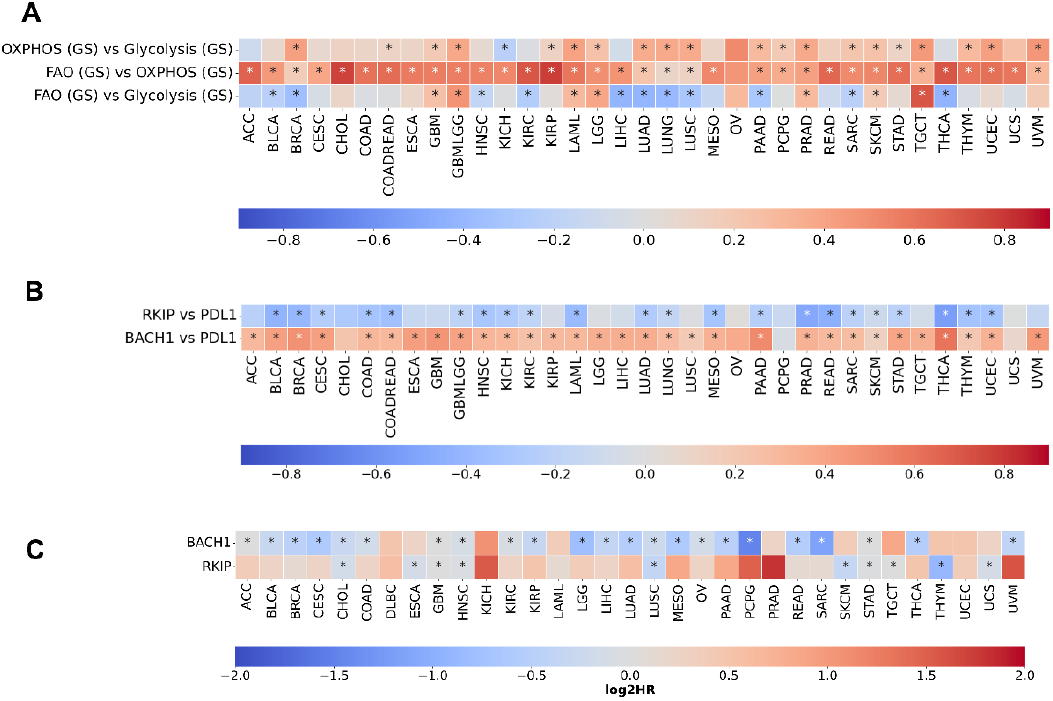
Correlations of RKIP and BACH1 expression with Immune Pathway signatures and Survival in TCGA. Heatmaps representing pairwise Spearman’s correlation coefficient of **A)** ssGSEA scores of metabolic pathway gene signatures: Glycolysis,) Fatty Acid Oxidation (FAO), Oxidative Phosphorylation (OXPHOS) with each other. **B)** ssGSEA scores of PD-L1 expression with RKIP and BACH1 **C)** Heatmap depicting Log2 normalized Hazard Ratios for RKIP and BACH1 across cancers. *: p-value < 0.05.

**Figure S2:**
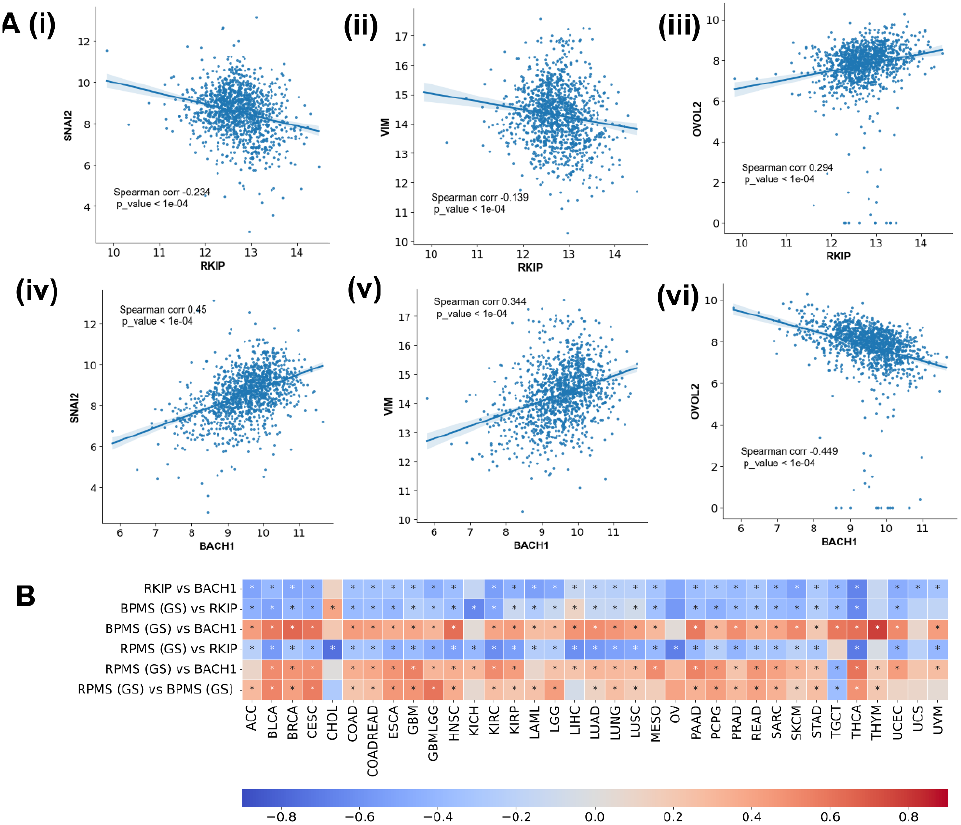
**A)** Scatter plots showing correlations of RKIP and BACH1 with SNAI2, VIM and OVOL2 in TCGA breast cancer data. **B)** Heatmap representing Spearman’s correlation coefficient of ssGSEA scores of RKIP Pathway Metastasis Signature (RPMS) and BACH1 Pathway Metastasis Signature (BPMS) with RKIP, BACH1 and each other, across various cancers. * denotes p-value < 0.05

**Figure S3:**
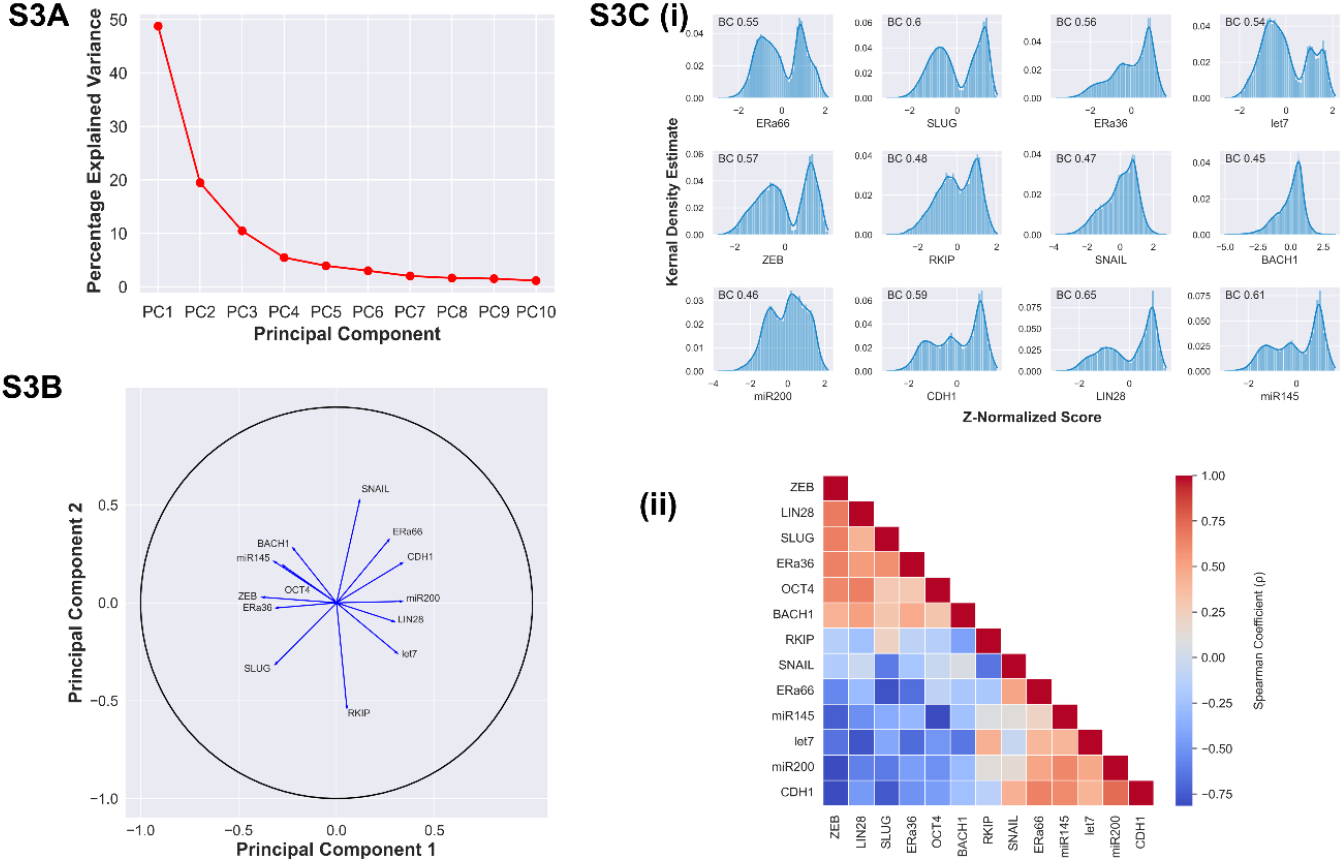
**A)** Scree-plot representing percentage explained variance captured by each principal component from PCA performed on steady state gene expression data obtained from RACIPE. **B)** PCA Correlation Circle representing correlations of RACIPE steady state gene data along PC1 and PC2 axes. **C)** i) Kernel Density Estimate plots of z-normalized expression values obtained from RACIPE simulation, for each node in the network (Figure 3A). Bimodality coefficients for each distribution are indicated as BC. ii) Pairwise correlation matrix showing Spearman correlation coefficient values of the corresponding pairs of genes. p-value < 10^−4^ for all pairwise correlations in the matrix.

**Figure S4:**
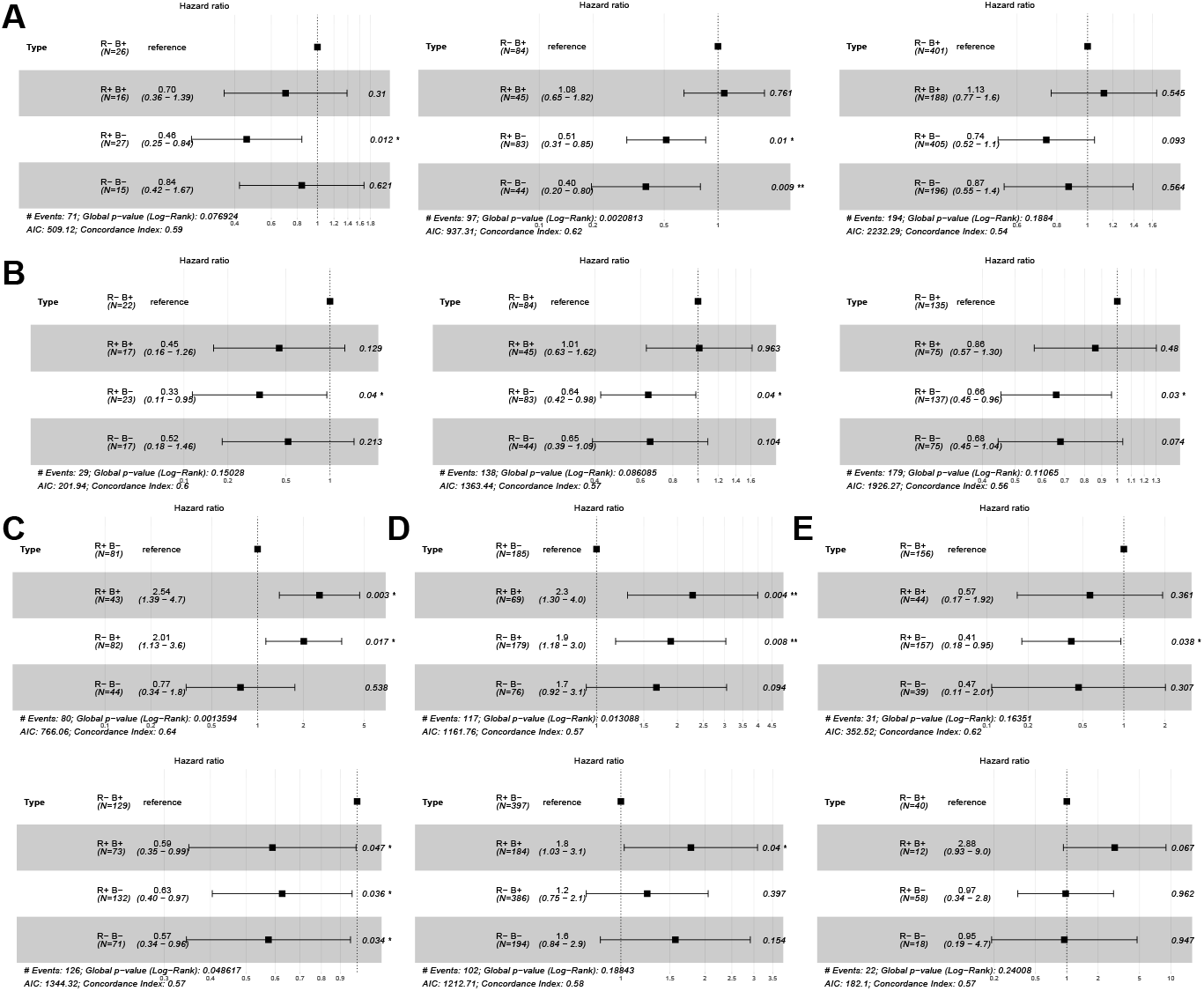
Forest plots comparing overall survival (OS) for different combinations of RKIP (low, high) and BACH1 (low, high) in TCGA samples. p-values based on log-rank test, and those with significant differences p < 0.05, p<0.01 and p<0.001 are marked with *, **, and ***, respectively. **A)** Hazard Ratios (HR) for Overall Survival in MESO (left), SARC (middle) and BRCA (right). **B)** HR for progression Free Survival in UVM (left), SARC (middle) and BLCA (right). **C)** HR for Disease-specific Survival in SARC (top) and BLCA (bottom). **D)** HR for disease-specific Survival in LGG (top) and BRCA (bottom). **E)** HR for disease-free Survival in THCA (top) and KIRC (bottom).

## Notes

### Competing Interest Statement

The authors have declared no competing interest.

